# Photoprotective demands predict external eye pigmentation in terrestrial mammals

**DOI:** 10.64898/2026.05.16.725635

**Authors:** Charlotte S. Streitferdt, Kai R. Caspar

**Affiliations:** Institute for Cell Biology, Heinrich Heine University Düsseldorf

**Keywords:** UV radiation, conjunctiva, peri-iridal tissues, coloration

## Abstract

The evolution of eye coloration in mammals and its potential ecological significance remain understudied. Evidence from anthropoid primates suggests that photoprotective demands are crucial determinants of pigmentation in the peri-iridal tissues, which encompass the conjunctiva and portions of the sclera peripheral to the iris. However, it is unclear to what extent these findings can be generalized. Here, we quantify peri-iridal brightness in a photographic sample of 62 terrestrial non-primate mammal species (*n* = 930). Phylogenetically-controlled analyses revealed significant effects of eye size as well as ecology on ocular pigmentation. Peri-iridal brightness exhibits a notable phylogenetic signal, correlates negatively with eye size and hence exposure to UV light, and is more pronounced in nocturnal species. Significant interspecific effects of latitude on peri-iridal brightness were not recovered, but tentative evidence for non-negligible impacts of this variable at the intraspecific level were found. Overall, these results align with and help to contextualize findings on primates and suggest that photoprotective demands importantly shape ocular appearance across the mammalian radiation. Furthermore, they have implications for hypotheses tying eye pigmentation chiefly to gaze signaling and provide a broad evolutionary framework for the emergence of human eye appearance.

## Introduction

The appearance of the mammalian eye is highly variable (Montiani-Ferreira et al., 2022). Important determinants of eye appearance include the shape of the palpebral fissure and pupil, as well as the coloration of the iris and the peri-iridal tissues surrounding it. Peri-iridal tissues are composed of the externally visible anterior portions of the sclera peripheral to the limbus and the overlying bulbar conjunctiva that adheres to it (Fig. 1A). In humans, these tissues give rise to the “white of the eye” (Perea-García et al., 2025). While the healthy mammalian sclera is, *in vivo*, almost invariably white to grey, the appearance of the bulbar conjunctiva is more diverse, ranging from plain black,to more or less patchily distributed shades of brown, to fully transparent (Montiani-Ferreira et al., 2022). Its pigmentation depends on the density and activity of melanocytes, which reside in the conjunctival epithelium (Rohen, 1962; Jacobiec, 2016). The conjunctiva’s contribution to the appearance of the eye is thus substantial (Rohen, 1962), which is why the use of the erroneous term “sclera” in reference to peri-iridal tissues is discouraged (Caspar et al., 2023; Perea-García et al., 2025). Even if no macroscopic pigmentation of the conjunctiva is observable, the tissue typically still harbors melanocytes that produce certain quantities of melanin granules (Oriá et al., 2013; Jacobiec, 2016). The same is true for the sclera (Hu et al., 2020).

**Fig. 1:**
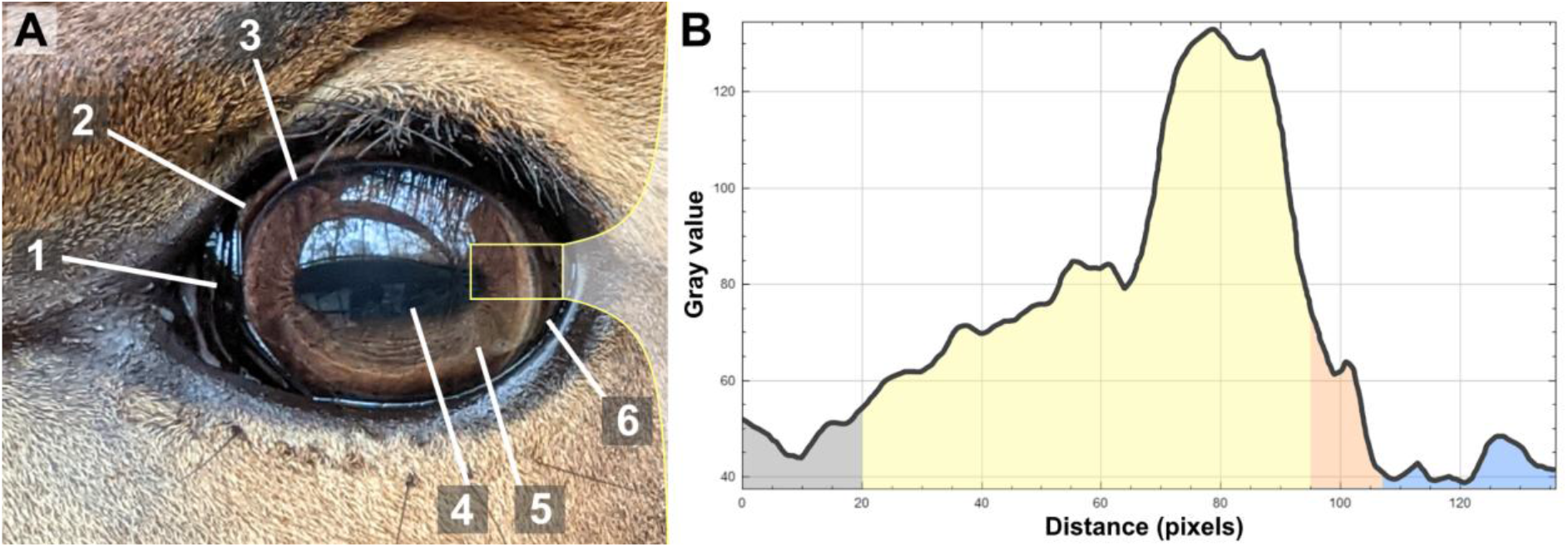
External eye appearance and quantification of peri-iridal brightness. **A**: Photo of the left eye of a bovid, exemplifying structures of the external eye in mammals. 1: Nictitating membrane, a conjunctival fold in the medial canthus of the eye. 2: Nasal aspect of the peri-iridal eyeball (sclera and overlying conjunctiva peripheral to the iris and cornea). Note comparatively light pigmentation. 3: Limbus, the anatomical interface between the cornea and the peri-iridal tissues. In many mammals, including the antelope shown here, a light circumferential ring forms in the limbal region. Due to its superficial similarity to the arcus senilis in humans, this understudied structure is at times referred to by the same name (e.g., Suedmeyer, 2006; Perea-García et al. 2021)). 4: Pupil. 5: Iris. 6: Temporal aspect of the peri-iridal eyeball. The brightness of this region of the eyeball was quantified. **B**: Gray value profile output of the ROI selected in **A** (yellow rectangle) as shown in ImageJ. Color shading indicates corresponding eye regions, with the pupil in grey, iris in yellow, limbus with “arcus senilis” in orange, peri-iridal tissues in blue. The highest and lowest gray scale values corresponding to the peri-iridal tissues were extracted for analyses.

In the last two decades, the significance of external eye appearance in mammals has been discussed with a strong focus on primates, especially humans (Perea-García et al., 2025). Most prominently, it was suggested that the coloration and salience of peri-iridal tissues primarily reflects a species’ use of eye gaze cues in social communication (Morris, 1985; Kobayashi & Koshima, 2001; Tomasello et al., 2007; Kano, 2023). However, accumulating evidence challenges this notion (reviewed by Perea-García et al., 2025) and other hypotheses have shifted into focus to explain the considerable spectrum of ocular appearances among primates but also other mammals (Caspar et al., 2021, 2023; Perea-García et al., 2021, 2025; Wacewicz et al., 2022). Among those is the idea that photoprotective demands are an adaptive driver of differences in peri-iridal pigmentation (Perea-García et al., 2022). We will refer to this here as the photoprotection hypothesis. Prolonged UV exposure can critically damage external ocular tissues and the conjunctival and limbal stem cells that reside in them (Grieve et al., 2015; Ramos et al., 2015), driving the emergence of debilitating pathologies such as certain eye cancers, keratopathies, and pterygium (MacKenzie et al., 1992; Gazzard et al., 2002; Yu et al., 2006; Yam & Kwok, 2014). Geographic latitude can act as a robust proxy for UV irradiance (Seckmeyer et al., 2008; Beckmann et al., 2014) and has been recovered as a significant interspecific predictor for ocular pigmentation in anthropoid primates: Species occurring at lower latitudes tend to show more strongly pigmented peri-iridal tissues (Perea-García et al., 2022; 2024). This pattern parallels findings for facial skin pigmentation in monkeys (Santana et al., 2012; 2013; Perea-García et al., 2024) and supports a crucial protective function of melanin. Intraspecifically, marked variation in conjunctival pigmentation and other photoprotective adaptations have been noted in human ethnicities originating from different latitudes (Mann, 1966; Kirschfeld, 1982; Jakobiec, 1984; Jakobiec, 2016). Outside of the primate order, evidence for such photoprotective effects on conjunctival pigmentation remains scarce. It should be noted that anthropoid primates are visual specialists exhibiting various striking ocular adaptations that are rare or absent among other mammal lineages (Kirk & Kay, 2004; Hall et al., 2012; Kelber & Jacobs, 2016). Despite differences in visual ecology, however, photoprotective specializations of ocular tissues should clearly be similarly beneficial to non-primate mammals, based on the pathophysiological principles outlined above.

Recently, Caspar et al.(2023) reported that the intensity of peri-iridal pigmentation was negatively correlated with the axial diameter of the eye in a small sample of placental mammals (*n*Taxa = 26). It can be observed that the palpebral fissure tends to become more elongated as mammalian eyes grow larger (although this has, so far, only been quantified in primates, to the best of our knowledge - Kobayashi & Kohshima, 2001). At the same time, at least in primates, large-bodied species with bigger eyes rely more on eyeball rather than head movements to visually scan their environment (Kobayashi & Kohshima, 2001). Both phenomena result in a greater exposure of the peri-iridal tissues to environmental UV radiation. The finding that an increase in eye size is associated with greater levels of melanin pigmentation is thus in line with predictions of the photoprotection hypothesis (Caspar et al., 2023; Perea-García et al., 2025). However, Caspar et al. (2023) sampled both nasal and temporal portions of peri-iridal tissues in an unbalanced fashion, which has since been shown to be a potential source of bias (Perea-García et al., 2024). Apart from geography and eye size, a species’ habitat use and lifestyle will affect peri-iridal UV exposure. For instance, UV irradiation is substantially lower for nocturnal species when compared to diurnal or cathemeral ones (see e.g., Seckmeyer et al., 2008). Therefore, according to the photoprotective hypothesis, one would expect lower peri-iridal pigmentation levels in species that are primarily active at night.

Here, we attempt to test the photoprotection hypothesis of peri-iridal pigmentation within a diverse sample of terrestrial non-primate mammals. A major limitation for studies on vertebrate eye pigmentation is the scarceness of eyeball specimens for comparative approaches. This results in a reliance on non-standardized digital photographs of live animals, on which the majority of recent studies on the topic are based on (e.g., Mearing et al., 2022; Perea-García et al. 2022; 2024; Caspar et al., 2021, 2023; Duran et al., 2024; Tabin & Chiasson, 2024). However, empirical evidence suggests that adequate sampling can sufficiently ameliorate noise from heterogeneous lighting conditions, making such photographs valuable for comparative biological research (Laitly et al., 2021).

Based on the patterns reported for primates and the predictions of the photoprotection hypothesis (Perea-García et al. 2022; 2024), we expected that large-eyed species and those occurring at lower latitudes would display increased levels of melanin pigmentation compared to small-eyed ones and those inhabiting high-latitudes. Additionally, we predicted that nocturnality would be negatively correlated with pigmentation levels due to decreased UV impact on peri-iridial tissues. For two species (African elephant - *Loxodonta africana*; lion - *Panthera leo*), we also explored potential intraspecific effects of latitude on peri-iridal pigmentation.

## Materials & Methods

### Sample composition

We quantified peri-iridall brightness in 62 therian mammalian species, following the taxonomy of the Mammal Diversity Database (Mammal Diversity Database, 2026). Only individuals identified as adults were considered in this study, as available evidence suggests that juvenile mammals tend to express weaker conjunctival pigmentation compared to mature ones (e.g., de Oliveira Garcia et al., 2021; Perea-García et al., 2024). Animals with recognizably aberrant pigmentation, such as melanistic or leucistic individuals, were also omitted. Domesticated forms were, for the most part, excluded, since their (ancestral) geographic ranges often cannot be meaningfully defined. Moreover, domesticated forms may display pronounced variation in ocular pigmentation and eyeball size across breeds (Howland et al., 2004; Caspar et al., 2023). Hence the only domesticated individuals covered were reindeer (*Rangifer tarandus*), for which the aforementioned issues were considered negligible. Primates were not sampled, given that extensive studies on this group are already available (Perea-García et al., 2022; 2024). Marine mammals were also not included.

We collect fifteen pictures per species, all publicly accessible online, based on sample size recommendations by Laitly et al. (2021) for using opportunistically taken, non-standardized photographs in comparative research on animal coloration. To qualify, the photograph had to allow differentiating between the peri-iridal tissues, iris, pupil and limbus. Photos that were visibly edited or noticeably over- or under-exposure were excluded. Photos showing animals with neutral facial expressions were preferably chosen to avoid sampling parts of the eyeball that are seldom exposed but might be visible during situations of heightened arousal (see e.g., Reefman et al., 2009). For each photo, only a single eye was sampled. If both eyes were visible, the better illuminated one was selected. The dataset is available in an OSF repository (https://doi.org/10.17605/OSF.IO/VMPYC).

Next to our main objective of testing predictions of the photoprotection hypothesis at the interspecific level, we also explored potential intraspecific effects of UV exposure on peri-iridal brightness at a smaller scale. It is commonly assumed that ocular tissues, including the iris and peri-iridal structures of the eye, do not tan in response to UV light. However, at least for the conjunctiva, this phenomenon has only received limited scientific attention so far (Perea-García et al., 2025) and data on non-human mammals are, to our best knowledge, absent. To consider the possibility of conjunctival tanning, we compared peri-iridal brightness in two species from our sample which exhibit strong conjunctival pigmentation: the lion and the African elephant. For these two species, sufficient numbers of suitable photos (*n* = 15) are available from both from their native low-latitude ranges as well as from individuals living in European and North American zoos (note as a caveat that since zoo animals are routinely transferred between institutions, the zoo in which an individual was photographed might not be the place it grew up or even spent the majority of its life in). If conjunctival pigmentation in these mammals is notably affected by tanning rather than guided by intrinsic factors, individuals from both species should display darker eyes within their native ranges compared to high-latitude zoo animals. We only used photos from the wild for those two species for the interspecific analyses.

### Quantification of peri-iridal brightness

Peri-iridal brightness was quantified as grayscale luminosity and measured using ImageJ (Schneider et al., 2012) by a single coder (CSS). For each photo, a rectangular region of interest (ROI) was defined, extending from the most temporally visible portion of the conjunctiva to the pupil (Fig. 1).To avoid confusion with the nictitating membrane and biases attributable to sampling different eyeball regions (see Perea-García et al., 2024) only the temporal aspect of peri-iridal tissues was analyzed (Fig. 1A). The ROI was selected in a way to cover the largest peri-iridal area possible while avoiding confounds such as reflections, shadows, and hair. The *plot profile* function was then applied to this ROI, generating a graph in which the image is subdivided into pixel columns, each yielding an average grayscale value (Fig. 1B). For each image, the minimum and maximum grayscale luminosity corresponding to the peri-iridal tissues (excluding the limbus) were extracted (Fig. 1B). The mean of these two values was incorporated into the analyses to act as a proxy for overall peri-iridal brightness.

Some previous work on primates (Perea-Garcia et al., 2022, 2024) used mean luminance values for a selected peri-iridal area to approximate tissue brightness (also using ImageJ software). These area estimates would initially appear more reliable than proxies derived from our approach, however, confounds such as conspicuous shadows and reflections often complicate the selection of a substantial peri-iridal area on available photos. To compare the agreement between the two methods and check the reliability of our approach, we coded images from 6 species with both and calculated a linear regression for the data (*n* = 90; species considered were *Cynictis penicillata, Didelphis virginiana, Hemigalus derbyanus, Macropus giganteus, Tapirus terrestris, Tragulus kanchil)*. For area measurements, we selected the largest continuous peri-iridal area of the visible eyeball that was free of aforementioned confounds. The coefficient of determination (adjusted R^2^) was found to be 0.91, pointing to a close correspondence of results between these approaches (Appendix, Fig. I). This corroborates the robustness of our method and suggests comparability of our findings with those of previous studies employing different methodologies.

Finally, we calculated intraclass correlation coefficients for peri-iridal brightness in the same six species, to ensure data coding reliability (*irr* package; Gamer et al., 2019). For all species, inter-rater agreement was good (*C. penicillata*: ICC = 0.77, *p* = 0.002) or excellent (*D. virginiana*: ICC = 0.91, *p* = 0.002; *H. derbyanus*: ICC = 0.96, *p* < 0.001; *M. giganteus*: ICC = 0.95, *p* < 0.001;*T. terrestris*: ICC = 0.97, *p* < 0.001; *T. kanchil*: ICC = 0.98, *p* < 0.001).

### Phylogenetic data and biological predictor variables incorporated into statistical models

To account for phylogenetic relatedness between species in our dataset, we used a consensus phylogeny calculated from 1000 trees downloaded from VertLife.org (Fig 1; Upham et al., 2019; see *Statistics* for details on phylogenetically-informed modelling).

Measurements of axial eye diameter (mm) were collated from various sources, especially Howland et al. (2004) and Hall et al. (2012) (see dataset for all references used). Latitudinal data was primarily derived from PanTHERIA (Jones et al., 2009; see dataset for exceptions). In line with other comparative studies on ocular pigmentation (Perea-García et al. 2022; 2024), we chose latitude as a proxy for UV irradiance (rather than, e.g., UV index). We selected a species’ range mid rather than minimum latitude after running models with either predictor and comparing DIC, which was marginally lower for the model incorporating the former (DIC = 9515.155 compared to DIC = 9515.553). Coding nocturnality was not straight-forward and can only represent an inexact approximation. We counted species as nocturnal when observations suggest primary activity at night and excluded those described as being cathemeral or crepuscular (note that crepuscularity in particular has been criticized as an inaccurate approximation of wild mammal activity patterns - Devarajan et al., 2025). We first consulted PanTHERIA for an initial assessment. We then compared those specifications with descriptions from accounts published in the journal *Mammalian Species*. In case of disagreement, we prioritized the reporting in *Mammalian Species*. When still in doubt, and for species not covered by either source, we took primary research articles into account. All respective references are included in the dataset. Predictor variables were not standardized.

### Statistics

#### All statistics were performed in R (RCore Team, 2024)

We analyzed factors influencing peri-iridal brightness within a phylogenetic framework, using a generalized linear mixed model with Markov chain Monte Carlo techniques (MCMCglmm) by aid of the package *MCMCglmm* (Hadfield, 2010). Fixed effects were assumed to have a significant impact on the response variable, peri-iridal brightness, when their 95% credible interval did not overlap with 0 (*α* = 0.05). We used the following model formula to study the effects of one categorial (nocturnality, yes/no) and two numeric predictors:

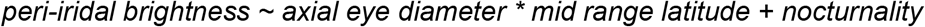

Phylogenetic relatedness and species affiliation were further incorporated as random effects. We ensured that the inclusion of these random effects increased our model’s fit compared to a null model by checking deviance information criteria (DIC) of model versions considering only one, both, or none of them (Appendix. Tab. 2).

We ran and validated our model following the documentation provided by Grieves et al. (2022) and fit it using an inverse gamma prior. We computed 15 × 10^6 model iterations, set the burn-in phase to 10000, and the thinning interval to 3000, resulting in a sample size of 4997.The function *raftery*.*diag()* from the *CODA* package (Plummer et al., 2006) was used to determine sufficient sample size. We ensured that autocorrelation was low by using the *autocorr*.*diag()* function from the same package and by viewing trace plots. The model was run three times to allow for a comparison of posterior mode and mean values and to calculate the Gelman–Rubin diagnostic (Gelman & Rubin, 1992). Results indicated that excellent convergence between models was achieved.

For the intraspecific comparison between high and low latitude populations of lions and African elephants, we ran a simple linear mixed effect model with species as a random effect, relying on the *lmer()* function from the lme4 package (Bates et al., 2015):

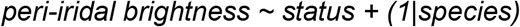

Peri-iridal brightness values were square root transformed to ensure normality of model residuals (checked by visual inspection of *qq* plots). The *lmerTest* package was used to assess the statistical significance of model coefficients (Kuznetsova et al., 2017). To further examine population differences at species level, we used two-sided Wilcoxon rank sum tests.

Intraspecific comparisons yielded results that were at least in parts compatible with the idea of latitudinal intraspecific effects on peri-iridal pigmentation. Hence, we checked if the inclusion of an additional random effect addressing this issue would increase the fit of our MCMCglmm. For this, we coded whether a sampled individual was photographed within (45.5 % of sample, *n* = 423) or outside (20 %, *n* = 186) its native latitudinal range in captivity. For 34.5 % (*n* = 321), locality information was insufficient, leaving the latitudinal status ambiguous. Inclusion of the aforementioned random effect (with three factor levels) marginally decreased the fit of the model (DIC = 9515.488) and it was found to only explain 0.03 % of random effect variance. Hence, we decided to not consider it for the MCMCglmm.

## Results

We found striking variation in peri-iridal brightness within our sample of 62 terrestrial mammal species (*n* = 930; Fig. 2). The brightest phenotypes were found among nocturnal marsupials and carnivorans, as well as lagomorphs-all species with small absolute eye sizes. The darkest peri-iridal tissues were observed in large-eyed ungulates. Peri-iridal brightness was significantly influenced by the eye’s axial diameter (Tab. 1, *p*MCMC = 0.0004). We found that eye size and peri-iridal brightness were inversely correlated within our species sample (Fig. 3). Thus, large eyes with greater exposure of the conjunctiva tended to show more intense peri-iridal pigmentation than smaller ones. Nocturnal species showed significantly brighter peri-iridal tissues and hence decreased pigmentation compared to taxa with different activity patterns (Tab. 1, *p*MCMC = 0.017). We did not recover an effect of latitude on peri-iridal brightness (Tab. 1, *p*MCMC = 0.297). However, we found a weak positive interaction between ocular diameter and the median latitudinal range of a species, the credible interval of which only marginally includes 0 (Tab. 1, *p*MCMC = 0.066). Hence, the aforementioned positive effect of eye size on peri-iridal pigmentation tends to be stronger in species occurring closer to the equator, although to a very limited extent.

**Fig. 2:**
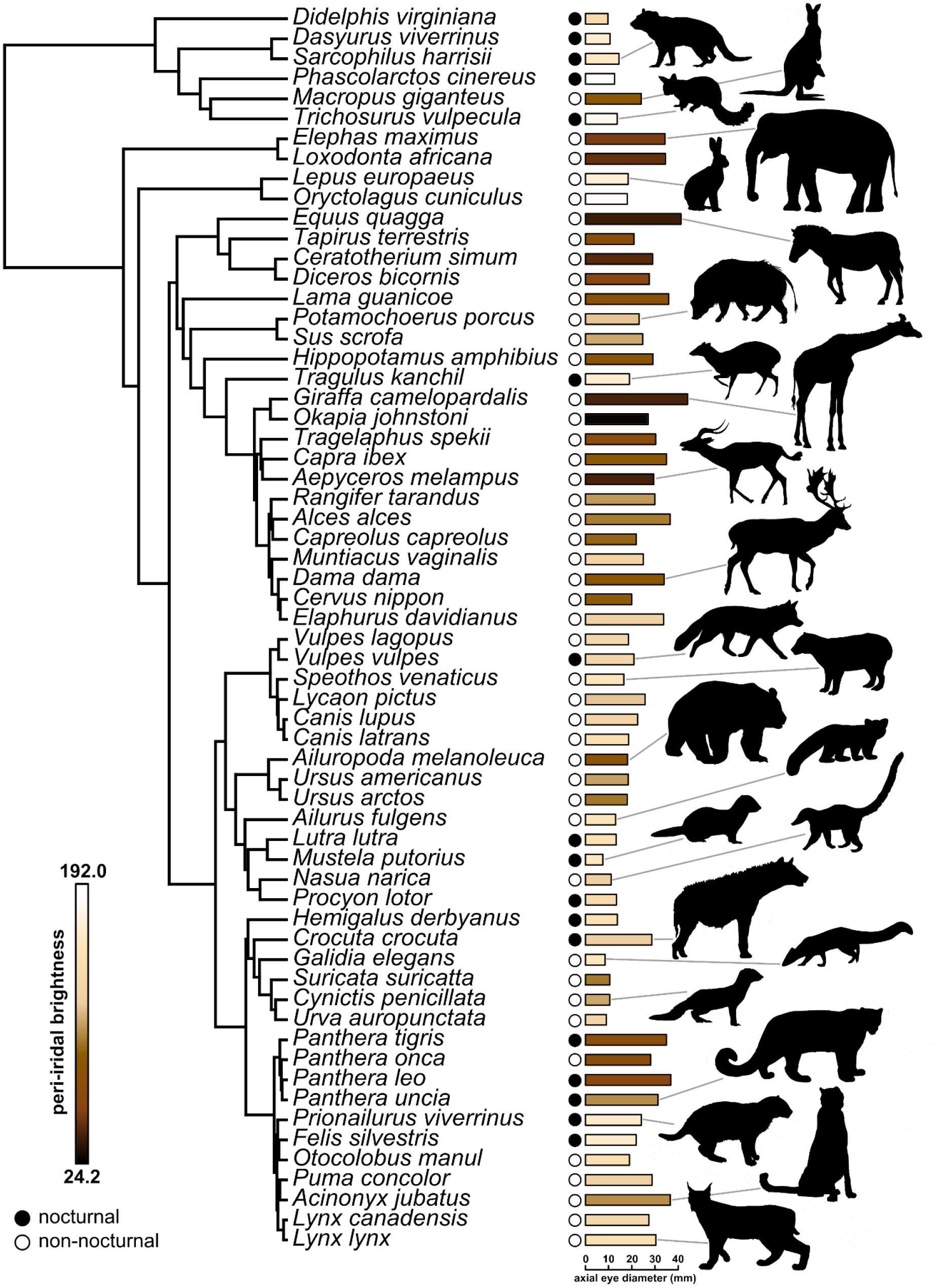
Peri-iridal brightness (grey scale values, color code represents an approximation) and phylogenetic relationships in a sample of 62 terrestrial mammal species. Bar length represents axial eye diameter (mm), circles indicate activity patterns (nocturnal: filled; not nocturnal: open). Silhouettes derive from phylopic.org (Keesey, 2025) and are in the public domain.

**Tab. 1:** MCMCglmm results quantifying effects of ocular diameter, range, their interaction, and activity on peri-iridal brightness. Given the close agreement between the three computed models, we only communicate the results from a single one of them. CI: credible interval.

| **coefficient** | **posterior mean** | **lower 95% CI** | **upper 95% CI** | **effective sample size** | **pMCMC** |
| --- | --- | --- | --- | --- | --- |
| **intercept** | 187.5 | 125.6 | 251.7 | 4757 | **<0.0002** |
| **axial eye diameter** | -3.59 | -5.13 | -1.95 | 4997 | **0.0004** |
| **mid. latitude** | -0.83 | -2.29 | 0.74 | 4507 | 0.297 |
| **nocturnality (present)** | 25.55 | 6.25 | 46.47 | 4997 | **0.017** |
| **eye diameter : mid. latitude** | 0.05 | -0.0003 | 0.11 | 4997 | 0.066 |

**Fig. 3:**
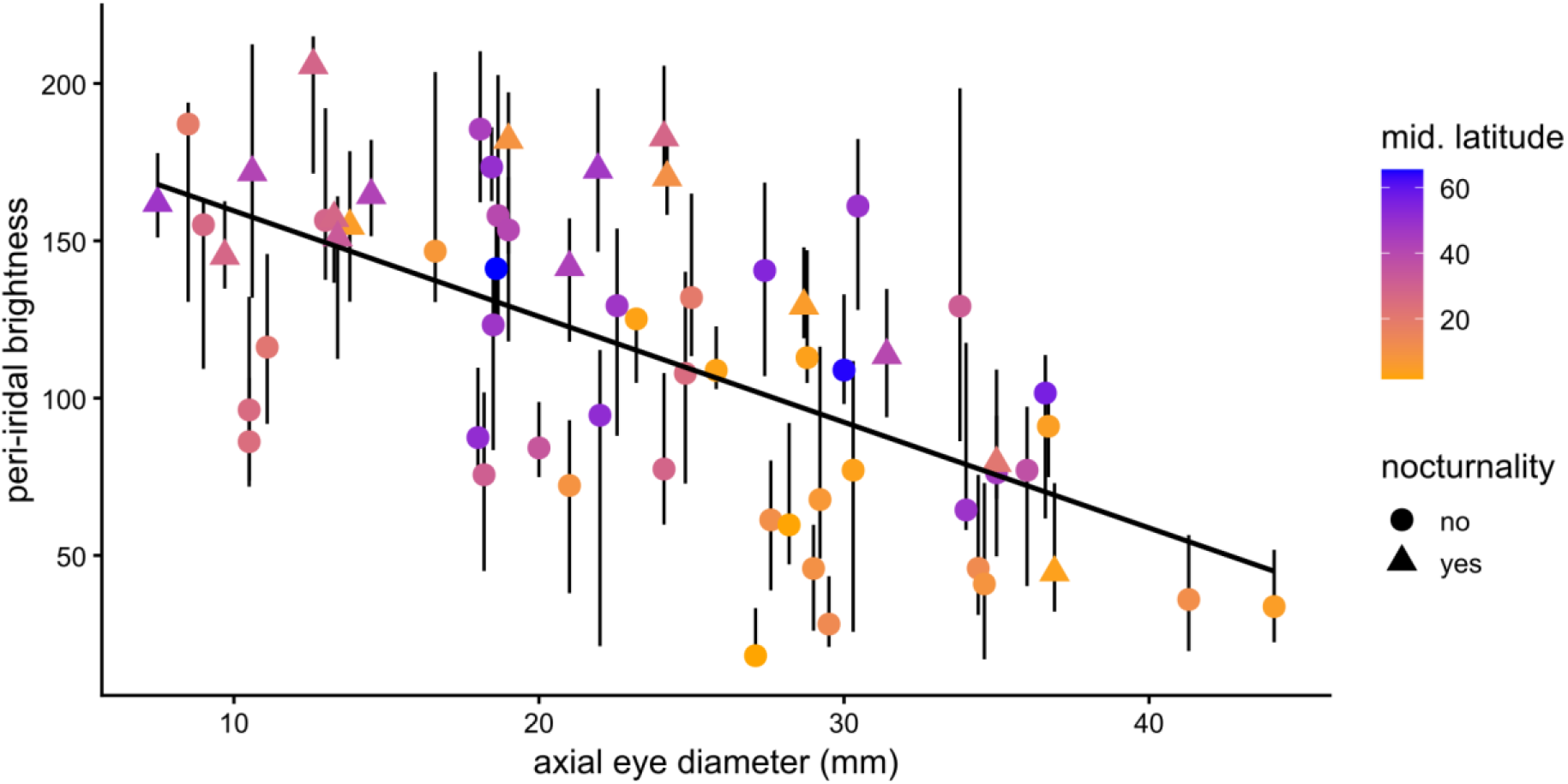
Peri-iridal brightness (grey scale values) is negatively correlated with axial eye diameter (mm) in 62 terrestrial mammal species. Median latitudes occupied by taxa are color-coded, activity patterns are represented by point shape. Point placement represents species’ median values, with whiskers indicating interquartile ranges.

Posterior means of random effect variance partitioning suggest a notable phylogenetic signal for peri-iridal brightness, with phylogeny explaining 28.8% of respective trait variation. Closely related species thus tend to show a similar expression of peri-iridal brightness. Additionally, 21.3% of variance was found to be species-specific, resulting in 49.9% residual variance. Details on random effects are provided in the appendix (Tab. II and Fig. II).

Finally, we performed an intraspecific comparison of peri-iridal brightness in wild and captive lions and African elephants, the latter living at notably higher latitudes in Europe and North America. In the interspecific dataset, we recovered significant differences between captive and wild individuals, with those from captivity presenting brighter peri-iridal tissues (Appendix, Tab. III, *p* = 0.032). When inspecting both species separately, differences between populations were significant in lions (*W* = 166, *p* = 0.03), but not in elephants (*W* = 131.5, *p* = 0.44; Fig. 4).

**Fig. 4:**
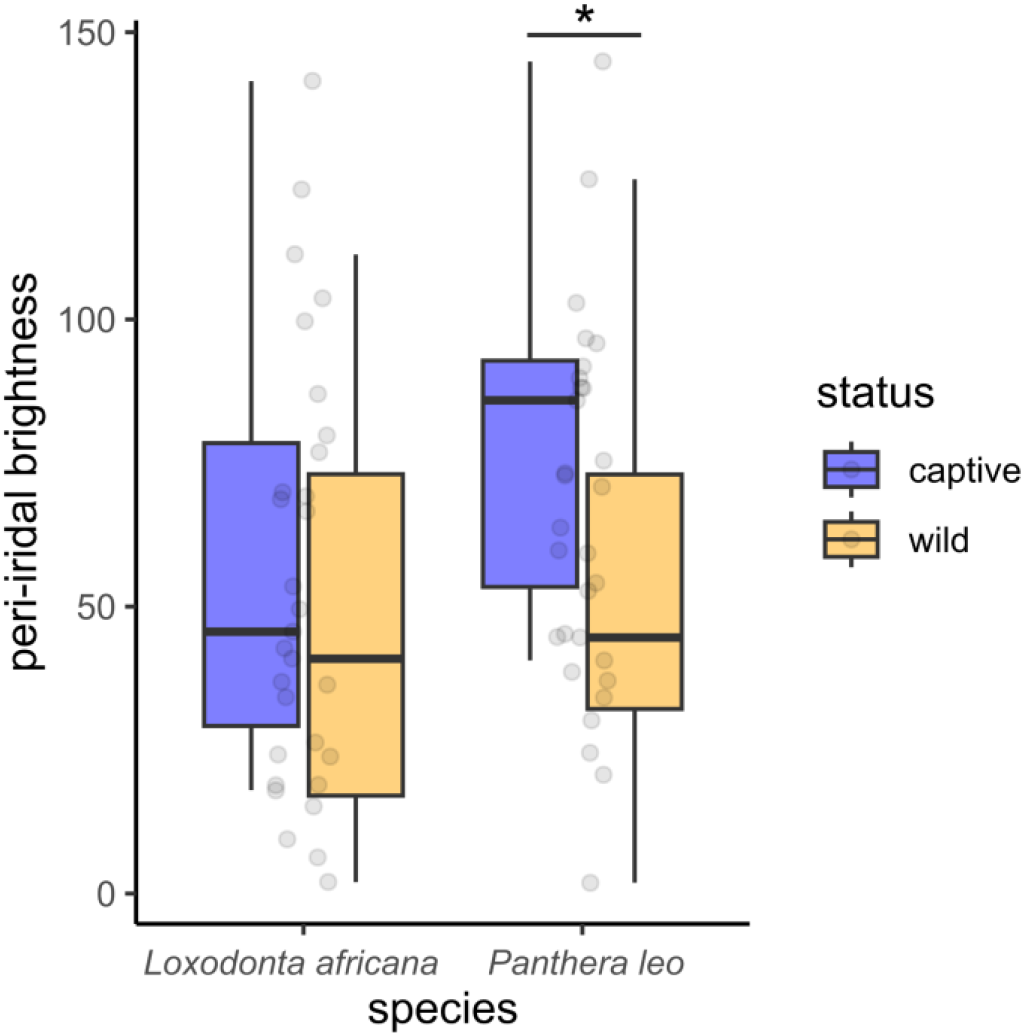
Peri-iridal brightness (grey scale values) in wild and captive populations of lions (*Panthera leo*) and African savannah elephants (*Loxodonta africana*). The difference between populations reaches statistical significance in lions (*p* = 0.03).

## Discussion

Patterns of peri-iridal pigmentation quantified for 62 terrestrial mammalian species overall align with predictions of the photoprotection hypothesis. These findings expand upon previous analyses on anthropoid primates (Perea-García et al. 2022; 2024; 2025) and suggest that photoprotective demands are important evolutionary drivers of external eye pigmentation across therian mammals. Our sample is biased towards large-bodied species and does not provide a balanced representation of therian taxa. Given limitations on the availability of suitable photos as well as eye ball measurements, this was unavoidable. However, our study provides testable predictions for future work on groups not covered in our analysis, such as rodents and bats.

We found a robust association between eye size and peri-iridal pigmentation (Tab. 1; Fig. 3), corroborating preliminary observations by Caspar et al. (2023). Thus, larger eyes that allow greater exposure of peri-iridal tissues to incoming light tend to exhibit darker pigmentation, and vice versa. Conspicuous deviations from this general trend might be indicative of marked intensification or relaxation of photoprotective demands, relating to a species’ ecology. One such example from our dataset is the meerkat (*Suricata suricatta*). This mongoose from south-western Africa has rather diminutive eyes (10.5 mm axial diameter) which nevertheless tend to be strongly pigmented. Its strict diurnality and unusual eye proportions represent convergences with anthropoid primates (Kirk, 2004). Meerkats inhabit arid open habitats (Van Staaden, 1994) exposing them to substantial UV radiation. Hence, their specific peri-iridal phenotype and lifestyle fit well with predictions of the photoprotection hypothesis. Interestingly, the dark facial mask typical of this species has been anecdotally suggested to reduce glare in the strongly lit environments that meerkats are found in (e.g., Badarnah, 2016). Whereas similar facial patterns are also known from various other carnivorans with different habitat preferences (Newman et al., 2005), such potential functional links between peri-iridal and facial coloration should be examined in future studies.

Activity patterns are important determinants of a species’ exposure to UV light, since UV irradiance is negligible at night (e.g., Seckmeyer et al., 2008). As predicted by the photoprotection hypothesis, we found that nocturnal mammals tend to exhibit lighter peri-iridal tissues than species with other activity rhythms (Tab. 1). While these findings are intuitive, their interpretation requires nuance since we coded activity as a binary variable. This decision was made for pragmatic reasons, but oversimplifies the actual spectrum of activity patterns encountered among the species included in our study. Furthermore, recent quantitative work based on global camera trap data has questioned the accuracy of traditional qualitative activity assessments, especially for ungulates and carnivorans (Devarajan et al., 2025). The same study also suggests greater intra-specific plasticity in circadian rhythms than is generally assumed for various mammal species. All this cautions the use of categorical predictors of diel activity, therefore calling for a considerate interpretation of our results.

Previous work on anthropoid primates suggests that latitudinal range, a proxy for UV irradiance, is a significant predictor of peri-iridal pigmentation (Perea-García et al. 2022; 2024). These findings are aligned with the photoprotection hypothesis but could not be replicated here for non-primate mammals. This was unexpected, especially given that the species studied herein cover a broader latitudinal range than previously examined anthropoids. Nevertheless, only a non-significant trend for a positive interaction between eye size and median latitude was found. Several explanations for this incongruence appear feasible. First, anthropoid primates tend to exhibit rather narrow latitudinal ranges, especially when compared to many of the ungulates and carnivorans represented in this study. This increases the informativeness of simple range proxies (e.g. median latitude) and might also result in less pronounced intraspecific variation in primate eye pigmentation. Second, anthropoids are almost exclusively diurnal (except for the genus *Aotus*) and ecologically as well as anatomically rather uniform. Hence, photoprotective demands would be expected to be higher and more homogeneous in this group compared to the non-primate mammals studied here. Habitat preferences and use have profound effects on the light irradiance a species encounters, potentially concealing latitudinal influences on UV exposure (see e.g., Dominy & Melin, 2020). For instance, animals living at high elevations and/or regions in which snow cover forms regularly may experience substantial UV exposure (Blumthaler et al., 1997; Chadyšiene & Girgždys, 2008) even at high latitudes. Coding these complex traits interacting with latitude for statistical approaches is challenging and beyond the scope of this study, but might provide the basis for important future insights on drivers of eye pigmentation.

Our analysis considers intraspecific variation in peri-iridal colouration, but we were unable to meaningfully link trait variability with latitudinal information, given often limited information about photo locations and our reliance on captive animals. While this also applies to other studies on eye pigmentation (e.g., Perea-García et al. 2022; Tabin & Chiasson, 2024) it confounds the detection of potential latitudinal effects. Whether conjunctival tanning, the hypothetical UV-induced acquired changes in conjunctival pigmentation, might contribute to phenotypic variation in mammals remains to be determined (see Perea-García et al. 2025). Results of our comparison of peri-iridal pigmentation in free-living and captive lions (but not elephants) photographed at low and high latitudes, respectively, are in line with the idea that such a phenomenon might exist and could influence macroscopic ocular phenotypes, at least in some species. However, meaningful conclusions clearly cannot be drawn from this. At most, these preliminary intraspecific observations call for careful decisions on photo sampling when assembling datasets such as ours and invite further investigations at the intraspecific level. In conclusion, whereas other lines of evidence suggest photoprotective demands to be drivers of peri-iridal pigmentation in terrestrial mammals, our analytical approach might not have been granular enough to detect the presence of latitudinal effects in our sample.

Although we deliberately did not analyse peri-iridal pigmentation in primates, our study does have implications for that animal group. Traditionally, interspecific differences in primate peri-iridal pigmentation have predominately been discussed in the context of social signalling, especially eye-gaze following (Morris, 1985; Kobayashi & Koshima, 2001; Tomasello et al., 2007; Kano, 2023). However, such hypotheses fall short of explaining the diversity of peri-iridal phenotypes in primates (Perea-García et al. 2025) and would also be unable to make sense of the patterns described here for other therian mammals. Our results suggest that photoprotection demands shape mammalian peri-iridal phenotypes and reinforce the idea that primates are also subject to such constraints. Hence, primatologists should consider abiotic factors more seriously as important determinants of eye coloration (Perea-García et al. 2022). For instance, callitrichids, the smallest of all monkeys, have been noted for exhibiting very bright eyes (Perea-García et al. 2022). It was argued that this phenotype results from a hypothetical self-domestication process and facilitates cooperative gaze signalling (Mearing et al., 2022). Yet, such an observation in small-eyed primates (ca. 12 mm axial eye diameter) fits well into the general mammalian pattern described here and could be explained by relaxed selection pressures on peri-iridal pigmentation. The assumption of a complex evolutionary scenario behind this ocular phenotype is thus rendered unnecessary (see also Caspar et al., 2023). On the other hand, humans present themselves as conspicuous outliers from the trends observed in other mammals. Although our species originates from low-latitude open habitats and has large eyes with a laterally elongated palpebral fissure, human peri-iridal pigmentation tends to be weakly developed (but note that non-pathologic intraspecific variability in this trait is higher than often assumed in the primatological literature - Perea-García et al. 2025). Hypotheses on why this might be the case have received extensive discussion recently (Wacewicz et al., 2022; Kano, 2023; Perea-García et al. 2025) and will not be reiterated here. We mention this example specifically to emphasize that photoprotective needs are not the only factor at play in the evolution of ocular pigmentation, regardless of whether we focus on primates or other mammals. For example, peri-iridal pigmentation is hypothesized to facilitate social communication in anthropoids primates in various ways (Perea-García et al. 2025) and there is evidence that the eyes are also of relevance for intraspecific communication in at least some carnivorans and ungulates (Somppi et al., 2014; Ueda et al., 2014; Wathan & McComb, 2014; see also Proctor & Carder, 2015, and Reefmann et al., 2009). Contrasting markings associated with the eyes are common in mammals, and interestingly also occur in lineages in which peri-iridal tissues are hardly visible, such as diurnal rodents (Le et al., 2026). This is despite the fact that most mammals exhibit substantially lower visual acuity than humans and other anthropoid primates (Veilleux & Kirk, 2014), making eye-mediated signalling more challenging. In some mammalian groups, peri-iridal pigmentation patterns might thus contribute to defining and amplifying facial expressions by making the eyeballs more salient (see Stringham, 2011, for relevant comments on bears).

## Conclusions and outlook

Mammalian eyes differ profoundly in the degree of peri-iridal pigmentation. The photoprotection hypothesis, originally formulated for primates, proposes that these patterns are importantly influenced by the UV irradiation of external ocular tissues. Our phylogenetic analysis of 62 therian mammals supports this assumption. We show that ocular diameter and nocturnal habits are significant predictors of peri-iridal brightness. Eye size, a proxy for UV exposure of peri-iridal tissues, is negatively correlated with brightness and pigmentation tends to be lower in species that are mostly active at night. Yet, an effect of latitude on peri-iridal brightness was unexpectedly not recovered, which might be explained by lifestyle and habitat characteristics concealing the influence of this variable within our ecologically diverse species sample.

So far, eye coloration in mammals has received surprisingly little attention, yet our results point to the presence of ecologically meaningful interspecific patterns that warrant further investigation. Future studies might focus more strongly on the extent to which pigmentation patterns are specific to certain taxa and reflect a species’ habitat structure and use. It will be of special importance in the long term to clarify how pigmentation patterns map onto the distribution of stem cells in the peri-iridal tissues, which also varies interspecifically (Grieve et al., 2015). This way, a comprehensive picture of the functional and adaptive relevance of peri-iridal pigmentation can eventually emerge.

## Acknowledgments

We thank Juan Olvido Perea-García for helpful discussions during the preparation of the manuscript and Teressa Wesley for constructive comments on an earlier draft of the text.

## Data availability

The data that support the findings of this study are openly available in OSF at https://doi.org/10.17605/OSF.IO/VMPYC.

## Author contributions

**Charlotte S. Streitferdt:** Investigation (lead); writing – original draft (equal); review and editing (equal); data curation (equal). **Kai R. Caspar**: Conceptualization (lead); investigation (supporting); methodology (lead); formal analysis (lead); writing – original draft (equal); writing – review and editing (equal); visualization (lead); data curation (equal).

## Appendix

**Tab. I:** Deviance information criteria (DIC) for models incorporating different random effects. Lower DIC values indicate better model fit, but ΔDIC of **≤** 2 are typically not considered meaningful (e.g., Spiegelhalter et al., 2022). The model incorporating species affiliation and phylogenetic relatedness was selected for the main analyses

| included random effects | DIC |
| --- | --- |
| species affiliation | 9515.1 |
| species affiliation, phylogenetic relatedness | 9515.2 |
| species affiliation, phylogenetic relatedness, native latitude | 9515.5 |
| phylogenetic relatedness | 9517.1 |
| none | 9851.1 |
| native latitude | 9851.5 |

**Tab. II:**
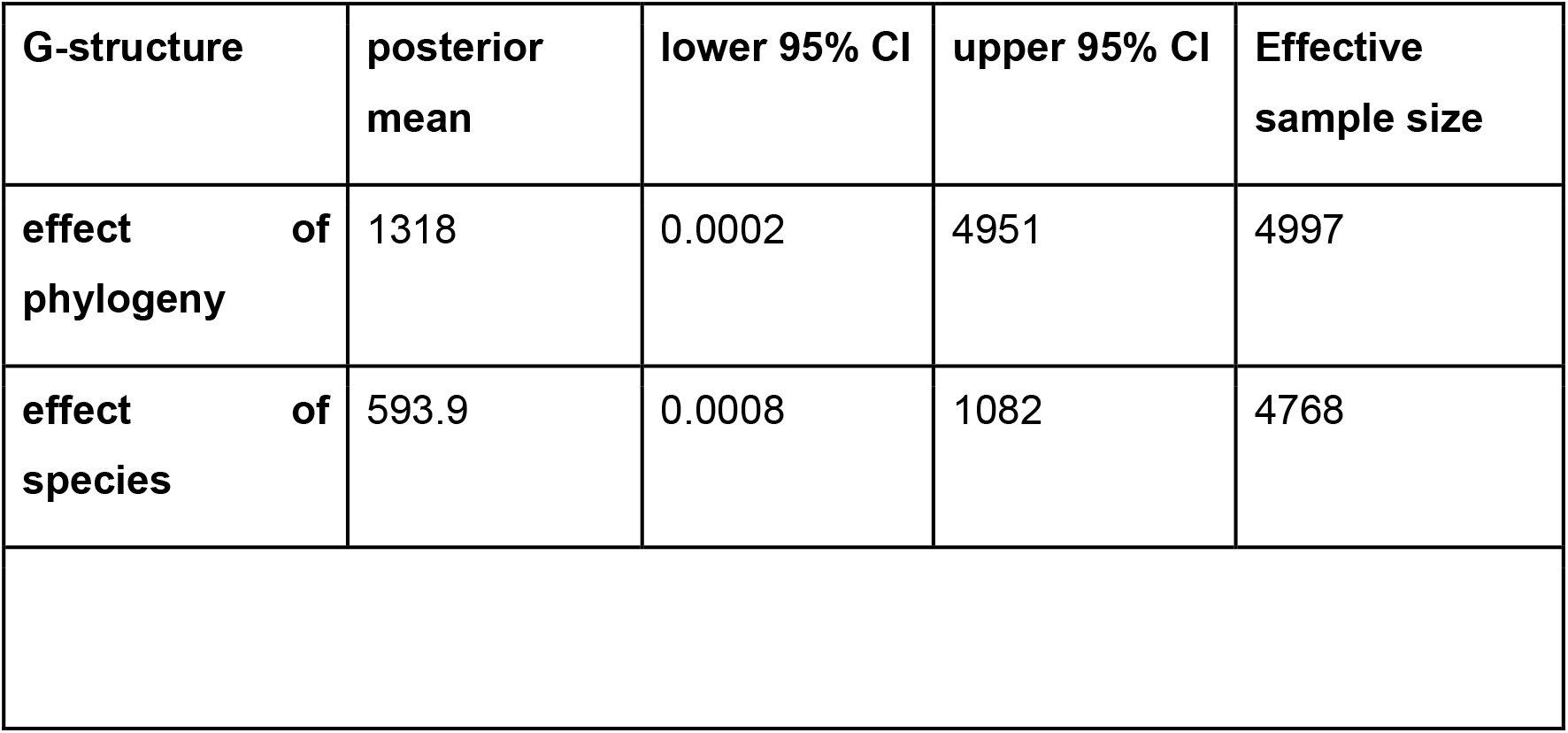
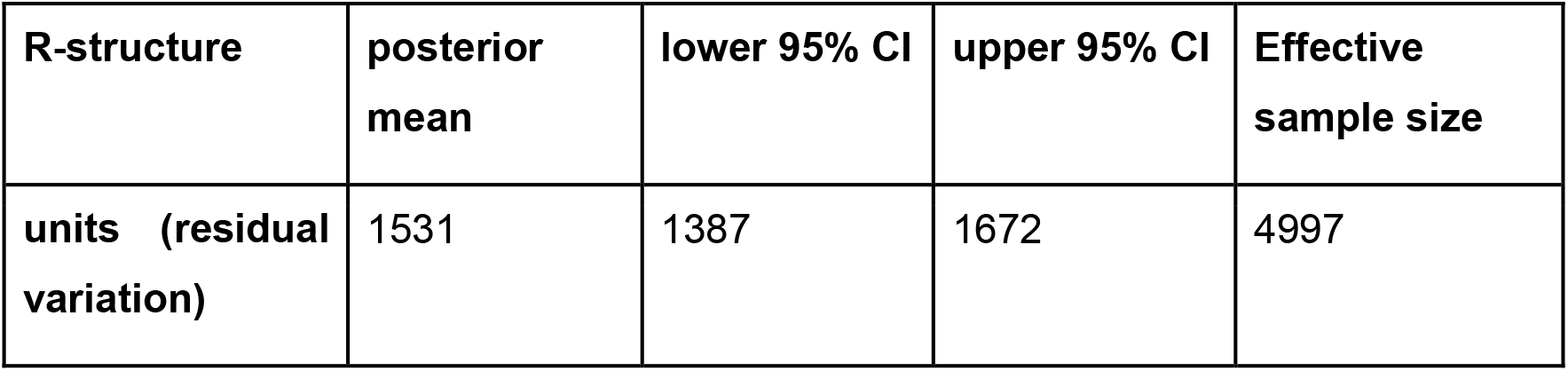
Random effects for MCMCglmm results quantifying effects of ocular diameter, range, their interaction, and activity on peri-iridal brightness. Given the close agreement between the three computed models, we only communicate the results from a single one of them, which is the same shown in Tab. 1 and discussed in the text. CI: credible interval.

| G-structure | posterior mean | lower 95% CI | upper 95% CI | Effective sample size |
| --- | --- | --- | --- | --- |
| effect of phylogeny | 1318 | 0.0002 | 4951 | 4997 |
| effect of species | 593.9 | 0.0008 | 1082 | 4768 |

| R-structure | posterior mean | lower 95% CI | upper 95% CI | Effective sample size |
| --- | --- | --- | --- | --- |
| units (residual variation) | 1531 | 1387 | 1672 | 4997 |

**Tab. III:**
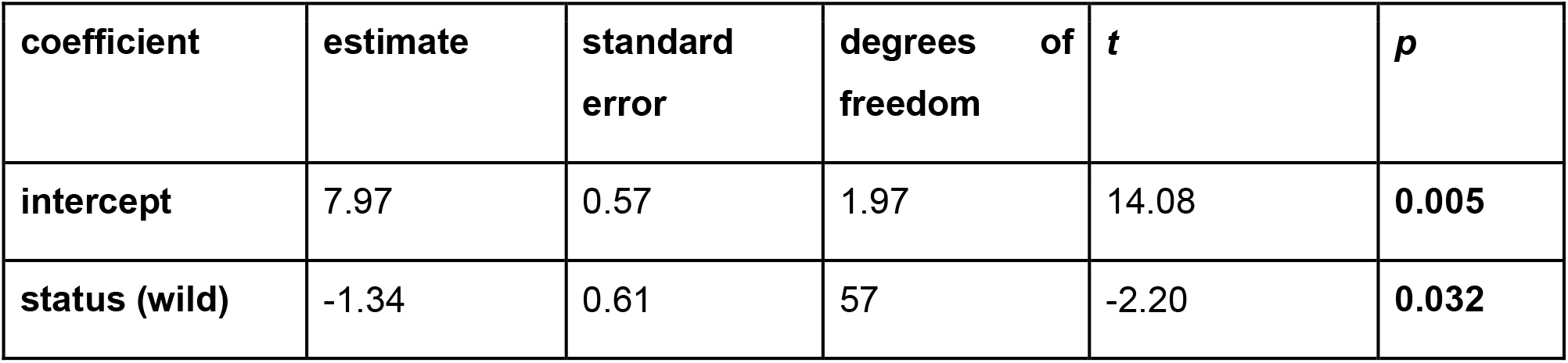
Results from a linear mixed effects model comparing peri-iridal brightness between captive and wild individuals of two species (African elephant, *Loxodonta africana*, and lion, *Panthera leo*).

**Fig. I:**
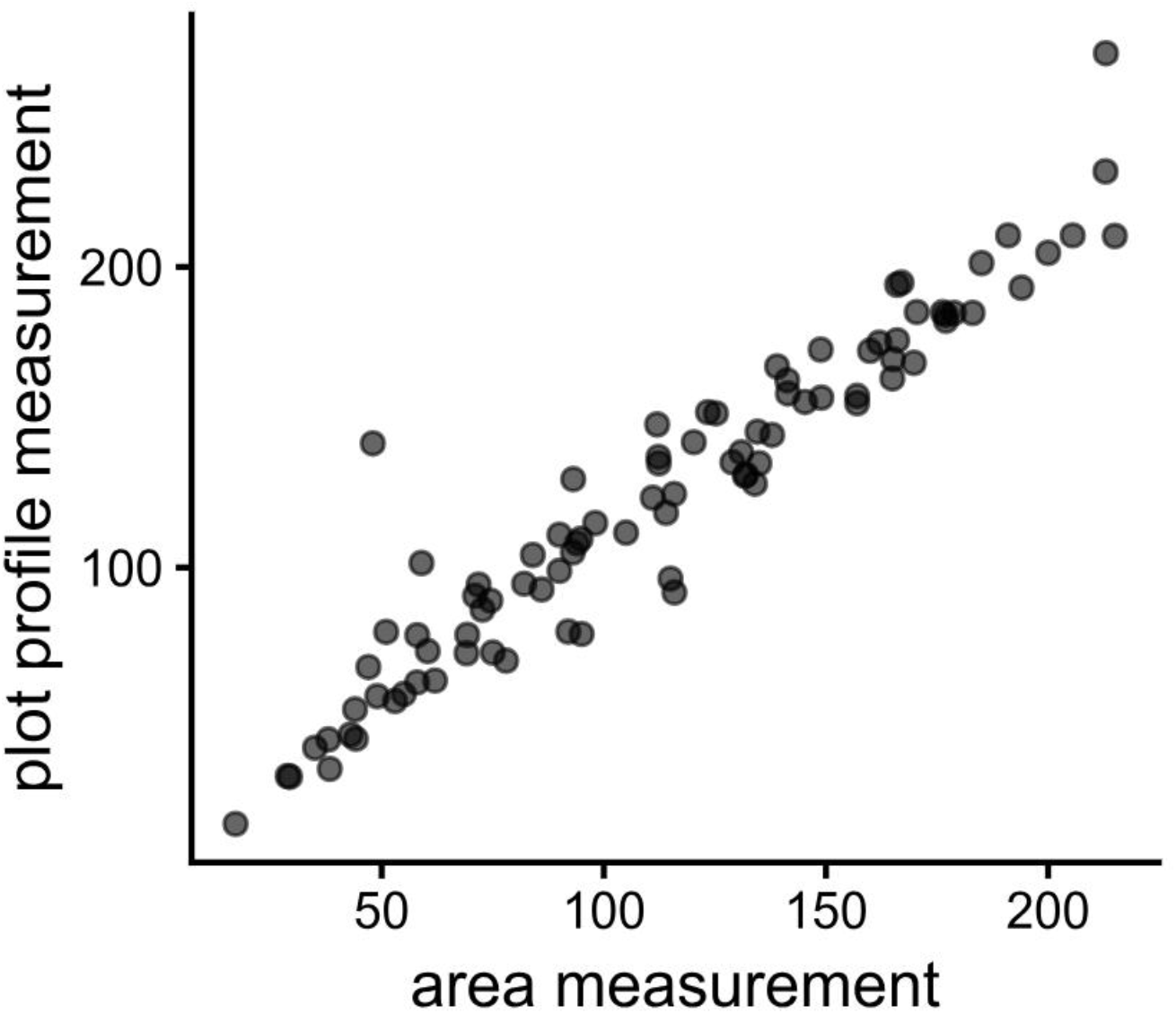
Correspondence of peri-iridal brightness measurements (derived from plot profiles of rectangular regions of interest and freely demarcated area selections), estimated in a subset of the study sample (*n* = 90). The coefficient of determination (adjusted R^2^) was found to be 0.91.

**Fig. II:**
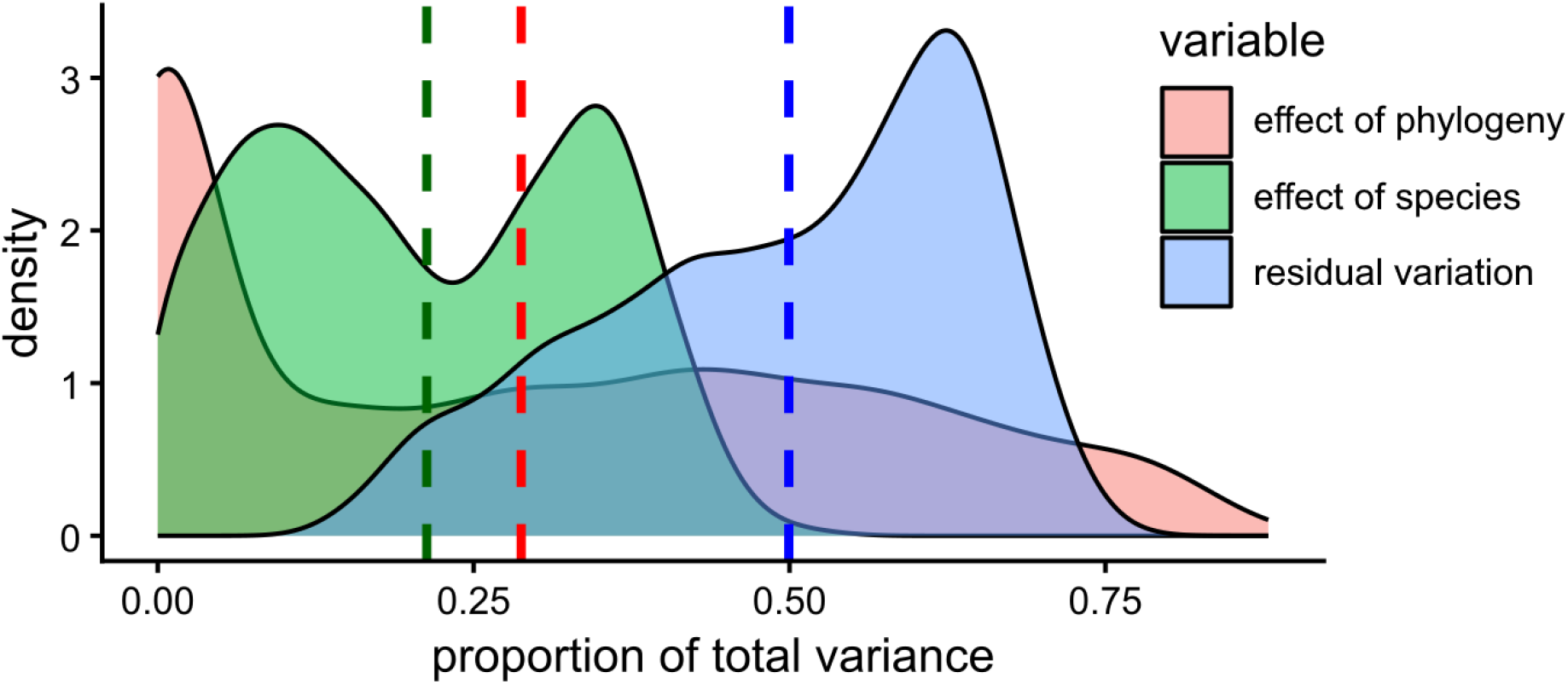
Variance partitioning of random effects from MCMCglmm. Dashed lines indicate posterior means of random effects. Red: effect of phylogeny, green: effect of species, blue: residual variation.

## References

Badarnah, L. (2016). Light management lessons from nature for building applications. Procedia Engineering, 145, 595–602.

Bates, D., Maechler, M., Bolker, B., Walker, S., Christensen, R. H. B., Singmann, H., Dai, B., Grothendieck, G., Green, P. & Bolker, M. B. (2015). Package ‘lme4’. Convergence 12(1), 2.

Beckmann, M., Václavík, T., Manceur, A. M., Šprtová, L., von Wehrden, H., Welk, E., & Cord, A. F. (2014). gl UV: a global UV-B radiation data set for macroecological studies. Methods in Ecology and Evolution, 5(4), 372–383.

Blumthaler, M., Ambach, W., & Ellinger, R. (1997). Increase in solar UV radiation with altitude. Journal of Photochemistry and Photobiology B: Biology, 39(2), 130–134.

Caspar, K. R., Biggemann, M., Geissmann, T. & Begall, S. (2021). Ocular pigmentation in humans, great apes, and gibbons is not suggestive of communicative functions. Scientific Reports 11(1), 12994.

Caspar, K. R., Hüttner, L., & Begall, S. (2023). Scleral appearance is not a correlate of domestication in mammals. Zoological Letters, 9(1), 12.

Chadyšiene, R., & Girgždys, A. (2008). Ultraviolet radiation albedo of natural surfaces. Journal of Environmental Engineering and Landscape Management, 16(2), 83–88.

de Oliveira Garcia, D., Estrela, G. C., Soares, R. T. G., Paulino Jr, D., Jorge, A. T., Rodrigues, M. A., … & dos Santos Honsho, C. (2021). A study on the morphoquantitative and cytological characteristics of the bulbar conjunctiva of the maned wolf (Chrysocyon brachyurus; Illiger, 1815). Anatomia, Histologia, Embryologia, 50(3), 439–447.

Devarajan, K., Fidino, M., Farris, Z. J., Adalsteinsson, S. A., Andrade-Ponce, G., Angstmann, J. L., … & de Oliveira Paschoal, A. M. (2025). When the wild things are: Defining mammalian diel activity and plasticity. Science Advances, 11(9), eado3843.

Dominy, N. J., & Melin, A. D. (2020). Liminal light and primate evolution. Annual Review of Anthropology, 49(1), 257–276.

Duran, E., Perea-García, J. O., Piepenbrock, D., Veefkind, C., Kret, M. E., & Massen, J. J. (2024). Preliminary evidence that eye appearance in parrots (Psittaciformes) co-varies with latitude and altitude. Scientific Reports, 14(1), 12859.

Gamer, M., Lemon, J., & Puspendra Singh, I. F. (2019). irr: various coefficients of interrater reliability and agreement. R package version 0.84.1. https://CRAN.R-project.org/package=irr

Gazzard, G., Saw, S. M., Farook, M., Koh, D., Widjaja, D., Chia, S. E. & Tan, D. T. H. (2002). Pterygium in Indonesia: prevalence, severity and risk factors. British Journal of Ophthalmology, 86(12), 1341–1346.

Gelman, A., & Rubin, D. B. (1992). Inference from iterative simulation using multiple sequences. Statistical Science, 7(4), 457–472.

Grieve, K., Ghoubay, D., Georgeon, C., Thouvenin, O., Bouheraoua, N., Paques, M. & Borderie, V. M. (2015). Three–dimensional structure of the mammalian limbal stem cell niche. Experimental Eye Research 140, 75–84.

Grieves, L. A., Gilles, M., Cuthill, I. C., Székely, T., MacDougall-Shackleton, E. A., & Caspers, B. A. (2022). Olfactory camouflage and communication in birds. Biological Reviews, 97(3), 1193–1209.

Hadfield, J. D. (2010). MCMC methods for multi-response generalized linear mixed models: the MCMCglmm R package. Journal of Statistical Software, 33(2), 1–22.

Hall, M. I., Kamilar, J. M., & Kirk, E. C. (2012). Eye shape and the nocturnal bottleneck of mammals. Proceedings of the Royal Society B: Biological Sciences, 279(1749), 4962–4968.

Howland, H. C., Merola, S., & Basarab, J. R. (2004). The allometry and scaling of the size of vertebrate eyes. Vision research, 44(17), 2043–2065.

Hu, D. N., Yao, S., Iacob, C. E., Giovinazzo, J., Rosen, R. B., Grossniklaus, H. E., & Sassoon, J. (2020). Quantitative study of human scleral melanocytes and their topographical distribution. Current Eye Research, 45(12), 1563–1571.

Jakobiec, F. A. (2016). Conjunctival primary acquired melanosis: is it time for a new terminology?. American Journal of Ophthalmology, 162, 3–19.

Jakobiec, F. A. (1984). The ultrastructure of conjunctival melanocytic tumors. Transactions of the American Ophthalmological Society, 82, 599–752.

Jones, K. E., Bielby, J., Cardillo, M., Fritz, S. A., O’Dell, J., Orme, C. D. L., … & Purvis, A. (2009). PanTHERIA: a species-level database of life history, ecology, and geography of extant and recently extinct mammals. Ecology, 90(9), 2648–2648.

Keesey, T. M. (2025). PhyloPic. https://www.phylopic.org

Kelber, A., & Jacobs, G. H. (2016). Evolution of color vision. In J. Kremers, R. Baraas, and N. Marshall (eds.) Human Color Vision. Springer Series in Vision Research, Vol. 5 (pp. 317–354). Springer, Cham.

Kirk, E. C., & Kay, R. F. (2004). The evolution of high visual acuity in the Anthropoidea. In F. C. Ross and R. F. Kay (eds.) Anthropoid Origins: New Visions (pp. 539–602). Springer, Boston.

Kirschfeld, K. (1982). Carotenoid pigments: their possible role in protecting against photooxidation in eyes and photoreceptor cells. Proceedings of the Royal Society of London. Series B. Biological Sciences, 216(1202), 71–85.

Kobayashi, H., & Kohshima, S. (2001). Unique morphology of the human eye and its adaptive meaning: comparative studies on external morphology of the primate eye. Journal of Human Evolution, 40(5), 419–435.

Kuznetsova, A., Brockhoff, P.B. & Christensen, R.H.B. (2017). lmerTest package: tests in linear mixed effects models. Journal of Statistical Software 82, 1–26.

Laitly, A., Callaghan, C. T., Delhey, K., & Cornwell, W. K. (2021). Is color data from citizen science photographs reliable for biodiversity research?. Ecology and Evolution, 11(9), 4071–4083.

Le, N. H. K., Li, S. H., Chiu, C. C., Weng, M. P., Chen, S. K., & Liao, B. Y. (2026). White eye-rings coevolved with diurnal behaviors as a trait enhancing visual appeal in rodents. Communications Biology. 10.1038/s42003-026-09916-0.

MacKenzie, F. D., Hirst, L. W., Battistutta, D. & Green, A. (1992). Risk analysis in the development of pterygia. Ophthalmology 99(7), 1056–1061

Mammal Diversity Database. (2026). Mammal Diversity Database (Version 2.0) [Data set]. Zenodo. 10.5281/zenodo.15007505

Mann, I. (1966). Culture, Race, Climate and Eye Disease: An Introduction to the Study of Geographic Ophthalmology. Charles C. Thomas, Springfield.

Mearing, A. S., Burkart, J. M., Dunn, J., Street, S. E., & Koops, K. (2022). The evolutionary drivers of primate scleral coloration. Scientific Reports, 12(1), 14119.

Montiani-Ferreira, F., Moore, B.A., Ben-Shlomo, G. (2022). Wild and Exotic Animal Ophthalmology – Vol. 2: Mammals. Springer, Cham.

Morris, D. (1985). Bodywatching: A Field Guide to the Human Species. Crown, New York.

Newman, C., Buesching, C. D., & Wolff, J. O. (2005). The function of facial masks in “midguild” carnivores. Oikos, 108(3), 623–633.

Oriá, A. P., Pinna, M. H., Almeida, D. S., da Silva, R. M. M., Pinheiro, A. C. O., Santana, F. O. & Oliveira, A. V. D. (2013). Conjunctival flora, Schirmer’s tear test, intraocular pressure, and conjunctival cytology in neotropical primates from Salvador, Brazil. Journal of Medical Primatology 42(6), 287–292.

Perea-García, J. O., Danel, D. P., & Monteiro, A. (2021). Diversity in primate external eye morphology: Previously undescribed traits and their potential adaptive value. Symmetry, 13(7), 1270.

Perea-García, J. O., Ramarajan, K., Kret, M. E., Hobaiter, C., & Monteiro, A. (2022). Ecological factors are likely drivers of eye shape and colour pattern variations across anthropoid primates. Scientific Reports, 12(1), 17240.

Perea-García, J. O., Ostner, J., Schülke, O., Kaburu, S., Majolo, B., Maréchal, L., … & Monteiro, A. (2024). Photoregulatory functions drive variation in eye coloration across macaque species. Scientific Reports, 14(1), 29115.

Perea-García, J. O., Teuben, A., & Caspar, K. R. (2025). Look past the cooperative eye hypothesis: reconsidering the evolution of human eye appearance. Biological Reviews, 100(5), 2038–2054.

Plummer, M., Best, N., Cowles, K., & Vines, K. (2006). CODA: Convergence diagnosis and output analysis for MCMC. R News, 6(1), 7–11

Proctor, H. S., & Carder, G. (2015). Measuring positive emotions in cows: Do visible eye whites tell us anything?. Physiology & Behavior, 147, 1–6.

Ramos, T., Scott, D. & Ahmad, S. (2015). An update on ocular surface epithelial stem cells: cornea and conjunctiva. Stem Cells International, 2015, 601731.

R Core Team (2021). R: A language and environment for statistical computing. R Foundation for Statistical Computing, Vienna, Austria. https://www.R-project.org/.

Reefmann, N., Wechsler, B., & Gygax, L. (2009). Behavioural and physiological assessment of positive and negative emotion in sheep. Animal Behaviour, 78(3), 651–659.

Rohen, J. W. (1962). Sehorgan der Primaten. In H. Hofer, A. H. Schultz, & D. Starck (eds.),: Primatologia, Handbuch der Primatenkunde, Band II/1, Lieferung 6, pp. 1–210. S. Karger Basel & New York.

Santana, S. E., Lynch Alfaro, J., & Alfaro, M. E. (2012). Adaptive evolution of facial colour patterns in Neotropical primates. Proceedings of the Royal Society B: Biological Sciences, 279(1736), 2204–2211.

Santana, S. E., Alfaro, J. L., Noonan, A., & Alfaro, M. E. (2013). Adaptive response to sociality and ecology drives the diversification of facial colour patterns in catarrhines. Nature Communications, 4(1), 2765.

Schneider, C. A., Rasband, W. S., & Eliceiri, K. W. (2012). NIH Image to ImageJ: 25 years of image analysis. Nature Methods, 9(7), 671–675.

Seckmeyer, G., Pissulla, D., Glandorf, M., Henriques, D., Johnsen, B., Webb, A., … & Carvalho, F. (2008). Variability of UV irradiance in Europe. Photochemistry and Photobiology, 84(1), 172–179.

Somppi, S., Törnqvist, H., Hänninen, L., Krause, C. M., & Vainio, O. (2014). How dogs scan familiar and inverted faces: an eye movement study. Animal Cognition, 17(3), 793–803.

Stringham, S. F. (2011). Aggressive body language of bears and wildlife viewing: a response to Geist (2011). Human-Wildlife Interactions, 5(2), 177–191.

Suedmeyer K (2006) Special senses. In: M. E. Fowler & S. K. Mikota (eds.) Biology, Medicine and Surgery of Elephants (pp 399–407). Blackwell Publishing, Ames.

Tabin, J. A., & Chiasson, K. A. (2024). Evolutionary insights into Felidae iris color through ancestral state reconstruction. iScience, 27(10), 110903.

Tomasello, M., Hare, B., Lehmann, H., & Call, J. (2007). Reliance on head versus eyes in the gaze following of great apes and human infants: the cooperative eye hypothesis. Journal of Human Evolution, 52(3), 314–320.

Ueda, S., Kumagai, G., Otaki, Y., Yamaguchi, S., & Kohshima, S. (2014). A comparison of facial color pattern and gazing behavior in canid species suggests gaze communication in gray wolves (Canis lupus). PloS One, 9(6), e98217.

Upham, N. S., Esselstyn, J. A., & Jetz, W. (2019). Inferring the mammal tree: species-level sets of phylogenies for questions in ecology, evolution, and conservation. PLoS Biology, 17(12), e3000494.

Van Staaden, M. J. (1994). Suricata suricatta. Mammalian Species, 1994(483), 1–8.

Veilleux, C. C., & Kirk, E. C. (2014). Visual acuity in mammals: effects of eye size and ecology. Brain Behavior and Evolution, 83(1), 43–53.

Wacewicz, S., Perea-García, J. O., Lewandowski, Z., & Danel, D. P. (2022). The adaptive significance of human scleral brightness: an experimental study. Scientific Reports, 12(1), 20261.

Wathan, J., & McComb, K. (2014). The eyes and ears are visual indicators of attention in domestic horses. Current Biology, 24(15), R677–R679.

Yam, J. C., & Kwok, A. K. (2014). Ultraviolet light and ocular diseases. International Ophthalmology, 34(2), 383–400.

Yu, G. P., Hu, D. N. & McCormick, S. A. (2006). Latitude and incidence of ocular melanoma. Photochemistry and Photobiology 82(6), 1621–1626.

## References

Spiegelhalter, D. J., Best, N. G., Carlin, B. P., & Van Der Linde, A. (2002). Bayesian measures of model complexity and fit. Journal of the Royal Statistical Society: Series B, 64(4), 583–639.

